# GTestimate: Improving relative gene expression estimation in scRNA-seq using the Good-Turing estimator

**DOI:** 10.1101/2024.07.02.601501

**Authors:** Martin Fahrenberger, Christopher Esk, Arndt von Haeseler

## Abstract

**Background:** Single-cell RNA-seq suffers from unwanted technical variation between cells, caused by its complex experiments and shallow sequencing depths. Many conventional normalization methods try to remove this variation by calculating the relative gene expression per cell. However, their choice of the Maximum Likelihood estimator is not ideal for this application.

**Results:** We present *GTestimate*, a new normalization method based on the Good-Turing estimator, which improves upon conventional normalization methods by accounting for unobserved genes. To validate *GTestimate* we developed a novel cell targeted PCR-amplification approach (cta-seq), which enables ultra-deep sequencing of single cells. Based on this data we show that the Good-Turing estimator improves relative gene expression estimation and cell-cell distance estimation. Finally, we use *GTestimate*’s compatibility with Seurat workflows to explore three common example data-sets and show how it can improve downstream results.

**Conclusion:** By choosing a more suitable estimator for the relative gene expression per cell, we were able to improve scRNA-seq normalization, with potentially large implications for downstream results. *GTestimate* is available as an easy-to-use R-package and compatible with a variety of workflows, which should enable widespread adoption.

## Introduction

Single-cell RNA-seq (scRNA-seq) provides new insights into cell diversity, differentiation and disease [1, 2, 3]. These insights are enabled by affordable high-throughput methods for the parallel sequencing of thousands of cells [4, 5]. However, they require many experimental steps, whose efficiency differs between cells, leading to high variability in the number of mRNAs captured. Additionally, sequencing depths as low as 20,000 reads per cell [6] and the nature of parallel sequencing introduce stochastic variation [5, 7, 8]. After accounting for PCR-duplicates among reads, a median of ∼5,000 *UMIs/cell* (number of sequenced mRNA molecules per cell) with a range of ∼500-20,000 *UMIs/cell* is typical for a high quality sample (Fig. 1a). This high technical variation between cells results in a low signal-to-noise ratio, which makes data analysis challenging. During data processing (Fig. 1b) *global-scaling normalization* methods [8] such as e.g. Seurat’s *NormalizeData* [9], scran’s *computeSumFactors* [10, 11] or scanpy’s *normalize_total* [12] account for the variation in *UMIs/cell* by calculating a single scaling-factor (or size-factor) per cell. Despite its simplicity, this approach has been shown to outperform more complex methods [13].

**Figure 1:**
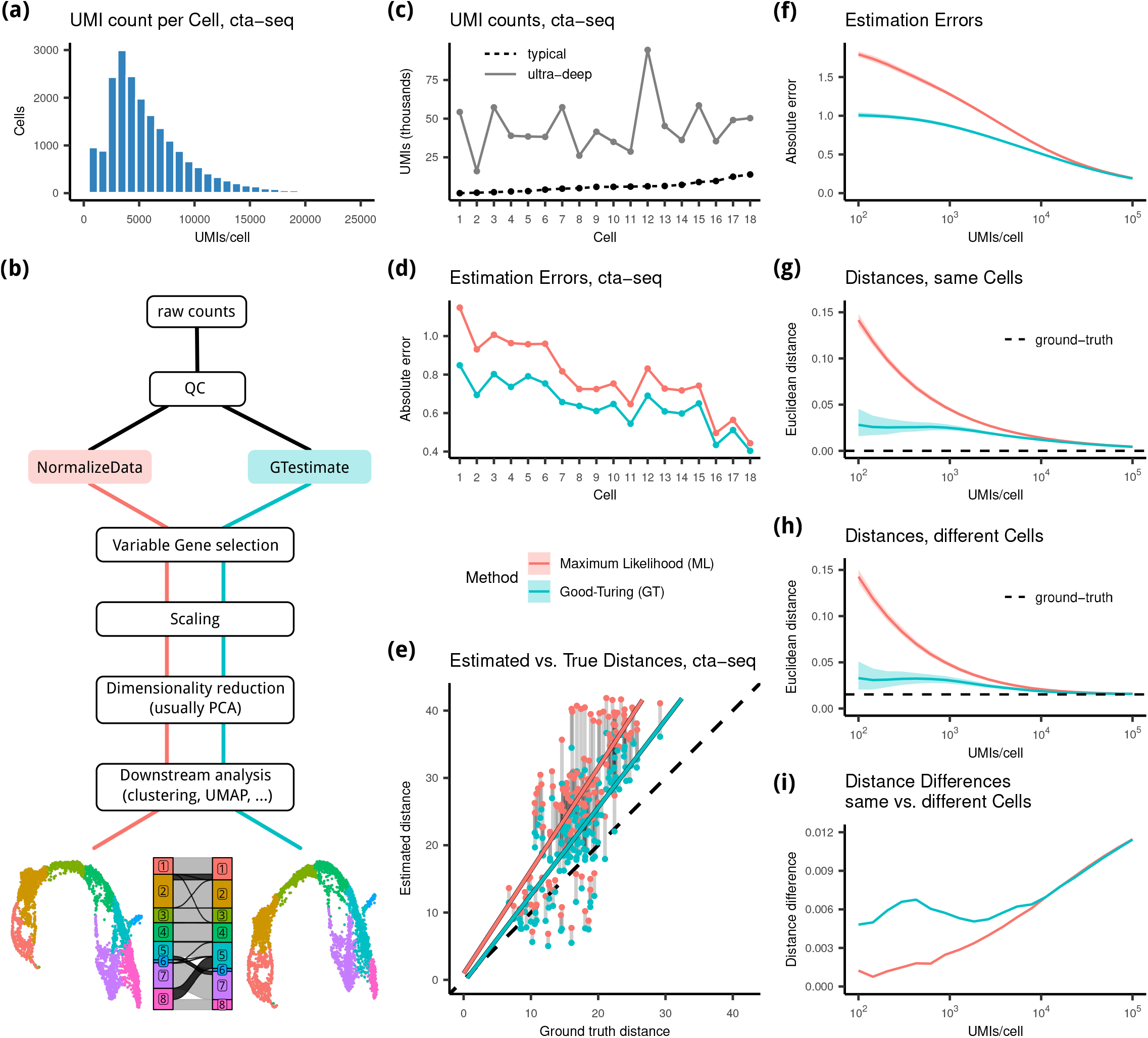
**(a)** Histogram of *UMIs/cell* for 17,653 cells in the cta-seq experiment before amplification. **(b)** Schema of a scRNA-seq analysis showing where *GTestimate* integrates into the workflow. **(c)** *UMIs/cell* for the 18 selected cells in the cta-seq experiment, before (*typical*) and after (*ultra-deep*) amplification. Cells ordered based on *UMIs/cell* in the *typical* cta-seq data. **(d)** Absolute error of the relative gene expression estimation in the cta-seq experiment. **(e)** Euclidean cell-cell distances in PCA-space in the cta-seq experiment. **(f)** Average absolute estimation error of the relative gene expression of a cell when subsampled to different *UMIs/cell*. **(g-h)** Mean Euclidean cell-cell distance in relative gene expression space, between two independent random samples of the same cell **(g)** between independent random samples of two different cells **(h). (i)** Difference between the mean cell-cell distances in **(g)** and **(h)**. Colored ribbons in **(f,g,h)** represent the 5% − 95% quantile range.

*Global-scaling normalization* inherently requires the calculation of the relative gene expression levels per cell. Although not typically discussed as such, the calculation used by these methods is a Maximum Likelihood estimation (ML) [14] of the relative gene expression frequency per cell.

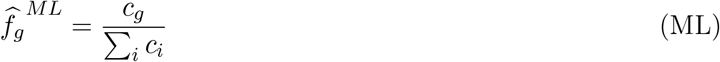

where *c* denotes the transcriptomic profile of the cell with a count *c*_*g*_ for each gene *g*.

However, at ∼5,000 *UMIs/cell* only ∼2.5% of the ∼200,000 mRNA transcripts in a typical mammalian cell [15] are sequenced and many expressed genes remain unobserved, as evident by the low *genes/cell* observed in scRNA-seq experiments (Suppl. Figure 1). ML then estimates the relative expression of unobserved genes as zero. This inherently leads to overestimation of the relative expression for observed genes, since the sum of all relative frequencies equals one 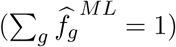.

To reduce this overestimation we propose a Simple Good-Turing estimator [16, 17].

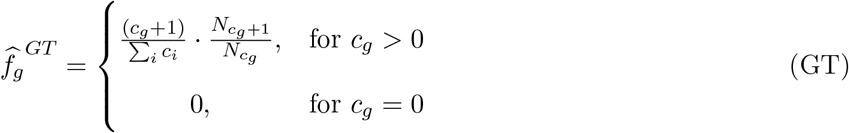

where 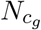 denotes the number of genes with count *c*_*g*_ in the cell.

GT adjusts the relative expression estimates of observed genes, particularly those with low counts, based on the frequency of each count value in the cell. This even enables an estimate for the relative expression of unobserved genes (for further details see Suppl. Info. 1.1).

In this study, we first compare the performance of GT and ML on novel ultra-deep sequencing data, and then show how GT improves downstream results, by integrating it into standard scRNA-seq analysis workflows. To achieve this we developed *GTestimate*, a new scRNA-seq normalization method centered around GT. *GTestimate* is an easy-to-use R-package designed to seamlessly replace Seurat’s *NormalizeData*.

## Results

### ultra-deep sequencing of single cells

Comparison between GT and ML requires ground-truth transcriptomic profiles of single cells. However, current simulation software cannot adequately emulate the complexity of scRNA-seq data and the choice of simulator may affect benchmarking results [18]. We therefore designed a cell targeted PCR-amplification strategy (cta-seq), which enabled us to sequence a small set of selected cells, from a *typical* sequencing run, a second time at a *ultra-deep* sequencing depth. This *ultra-deep* sequencing data contains an average of 23 million reads per cell, a stark contrast to the average 16,965 reads for the same cells in the *typical* data (Suppl. Figure 2). This led to a ∼7.4 fold increase in *UMIs/cell* (Fig. 1c) and a ∼3.3 fold increase in *genes/cell* (Suppl. Figure 3). We then used the relative gene expression levels of these *ultra-deep* profiles as the ground-truth for these cells.

### Performance of GT and ML

Based on the cta-seq data we then evaluated GT and ML. When we applied GT and ML to the *typical* profiles and compared the results to the ground-truth, GT consistently showed a lower estimation error across all 18 cells, by ∼17% on average (Fig. 1d).

Relative gene expression profiles are the basis of most scRNA-seq analysis (Fig. 1b), such as the calculation of cell-cell distances in PCA-space (often used as a measure for the similarity between two cells). We therefore also calculated cell-cell distances between the *typical* profiles, once based on GT and once based on ML, and compared the results to the cell-cell distances between the *ultra-deep* profiles. We observed a 36% reduction of the distance estimation error when using GT instead of ML (Fig. 1e, Suppl. Table 1).

Since *UMIs/cell* vary drastically (Fig. 1a) we further assessed the performance of GT and ML at different *UMIs/cell*. We applied GT and ML to random subsamples of the cell with the highest *UMIs/cell* in the *ultra-deep* cta-seq data (Cell 12, at 94,440 UMIs) and compared the estimates to the ground-truth expression profile of this cell. Similar to before (Fig. 1d) the estimation error for both GT and ML decreased with increasing *UMIs/cell* and GT consistently showed a lower error than ML, especially at low *UMIs/cell* (Fig. 1f).

Next, we assessed the impact of *UMIs/cell* on cell-cell distances. We first compared the mean distance between two random samples of the same cell (cell 12), both sampled to the same *UMIs/cell*. This distance was calculated in relative gene expression space and should approach zero for high *UMIs/cell*. However, ML led to grossly overestimated distances at small *UMIs/cell* (Fig. 1g). The estimated distance after ML additionally showed strong correlation to the *UMIs/cell*, which is problematic as we assume that most of the observed variation in *UMIs/cell* is technical noise. In contrast, GT did not show correlation to the *UMIs/cell* and demonstrated lower distance estimation errors overall.

We then examined the distances between two distinct cells by also drawing random samples from the cell with the second highest *UMIs/cell* in the *ultra-deep* cta-seq data (cell 15, at 58,589 UMIs), which is of a different cell-type. We calculated the distances between the sampled profiles of cell 12 and cell 15 at varying *UMIs/cell*. We again saw large overestimation of the distances when using ML, while using GT strongly reduced this error. For high *UMIs/cell* the estimated distances converged to the true distance of 0.015 (Fig. 1h).

When based on ML, the estimated distances between identical cells (Fig. 1g) and distinct cells (Fig. 1h) were almost the same for low *UMIs/cell*. This makes it very difficult to e.g. distinguish between cell-types. However, when we used GT as the basis for these distances we saw a much clearer separation between identical cells and cells of different cell-type, for cells with *<* 10, 000 *UMIs/cell* (Fig. 1i).

### *GTestimate*’s impact on downstream results

After showing GT’s advantages for relative gene expression estimation and cell-cell distance estimation, we examined how our GT based normalization method *GTestimate* impacts downstream results. The difference between *GTestimate* and other *global-scaling normalization* methods is only in the estimator used, all other settings can be adjusted to be equivalent to e.g. scran’s *computeSumFactors* or scanpy’s *normalize_total*. At default settings *GTestimate* behaves identically to *NormalizeData*, including the same log-transformation. We therefore used *NormalizeData*, as a representative of ML based *global-scaling normalizations* for all following comparisons. However, we would expect similar results when comparing to other *global-scaling normalization* methods.

We first assessed *GTestimate*’s impact on cell-type clustering by reanalyzing the pbmc3k data-set of peripheral blood mononuclear cells [19]. Here, normalization with *GTestimate* instead of *NormalizeData* resulted in 4.6% of cells being assigned to a different cluster (Fig. 2a), mostly among the Naive CD4 T-cells, Memory CD4 T-cells and CD8 T-cells.

**Figure 2:**
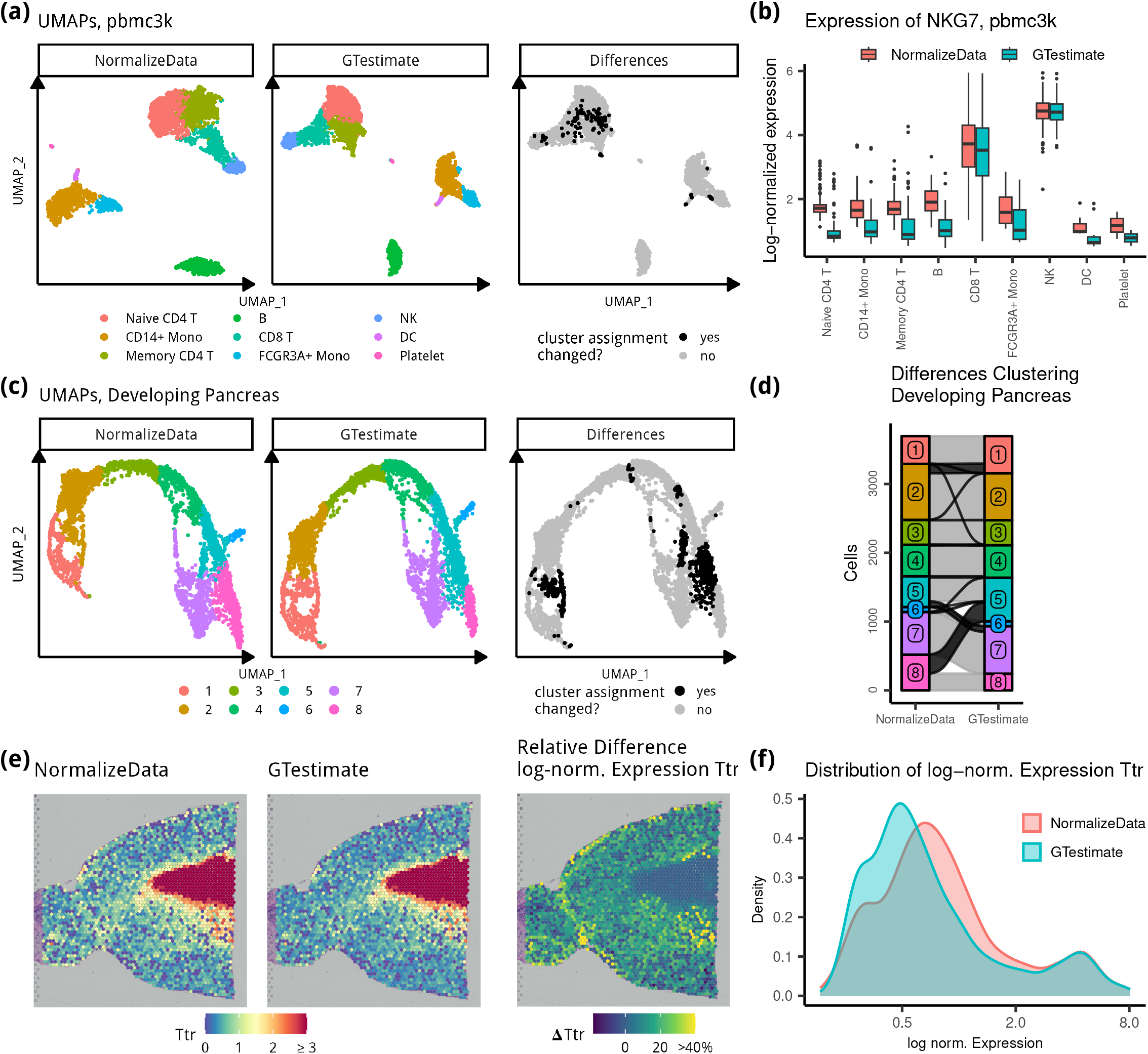
**pbmc3k: (a)** UMAPs based on *NormalizeData* and *GTestimate*, and UMAP highlighting differences in cluster assignment. **(b)** Boxplot showing log-normalized expression of *NKG7* per celltype (zeroes not shown). **Developing Pancreas: (c)** UMAPs based on *NormalizeData* and *GTestimate*, and UMAP highlighting differences in cluster assignment. **(d)** Sankey diagram showing the differences in cluster assignment based on *NormalizeData* and *GTestimate*. **Spatial Transcriptomics: (e)** log-normalized gene expression of *Ttr* based on *NormalizeData* and *GTestimate* as well as percent difference in log-normalized expression of *Ttr* between *NormalizeData* and *GTestimate*. **(f)** Density plot showing the distribution of log-normalized gene expression values of *Ttr* for *NormalizeData* and *GTestimate*.

We additionally analyzed a developing pancreas data-set [20], characterized by more gradual cell-type transitions compared to the pbmc3k data-set. After normalization with *GTestimate* instead of *NormalizeData*, 14.6% of cells were assigned to a different cluster (Fig. 2c,d).

While the correct classification of cells in both of these data-sets remains unknown, our results in Fig. 1 suggest that *GTestimate* provides a better basis for this classification.

To examine the impact of *GTestimate* on the expression estimates of individual genes we considered the log-normalized expression of cell-type specific marker genes in the pbmc3k data-set. As an example we used *NKG7* a highly specific NK-cell and *CD8* + T-cell marker [21]. When using *GTestimate* instead of *NormalizeData*, the log-normalized expression of *NKG7* remained constant in NK-cells and *CD8* + T-cells, but was reduced in all other cell-types (Fig. 2b). *GTestimate* therefore resulted in clearer separation of NK-cells and *CD8* + T-cells from other cell-types. We observed this for nearly all marker genes described in Seurat’s pbmc3k tutorial (Suppl. Figure 4). These differences may explain some of the observed changes in clustering.

Finally, we applied *GTestimate* to the spot-wise normalization of a Spatial Transcriptomics data-set of the mouse brain [22]. In this data-set, normalization with *GTestimate* and *NormalizeData* resulted in 17 and 19 clusters respectively (Suppl. Figure 5, Figure 6, Figure 7), we therefore refrained from any cluster based comparisons of *GTestimate* and *NormalizeData*. However, the spatial coordinates enabled examination of area specific marker genes, independent of the clustering. As an example we considered the log-normalized expression of the choroid plexus marker gene *Ttr* (Fig. 2e). When using *GTestimate* we saw a reduction of the unspecific expression of *Ttr* for spots outside the choroid plexus. Here, *GTestimate* showed up to 50% reduction of the log-normalized expression, compared to *NormalizeData*, while expression estimates inside the choroid plexus remained constant (Fig. 2e). This resulted in clearer separation of the choroid plexus spots from the surrounding tissue as shown by the distribution of expression values of *Ttr* (Fig. 2f).

When we additionally considered the *UMIs/spot* (Figure 8), we saw a negative correlation between the change in log-normalized expression of *Ttr* and *UMIs/spot*. This supports previous observations that *NormalizeData* overestimates the expression of *Ttr* in areas with low *UMIs/spot*. Whereas, *GTestimate* reduces this overestimation and improves the signal-to-noise ratio.

## Discussion

In summary, the estimation of relative gene expression is a central part of scRNA-seq data analysis, which has not received the same attention as other steps. We have shown that replacing the standard ML with GT improves relative gene expression estimation, without requiring expensive computations. By improving the signal-to-noise ratio at this basic level, our new normalization method *GTestimate* can have large impact on downstream results.

In the validation we avoided potential issues with simulated data by employing a novel cell targeted PCRamplification strategy to sequence the same cells at two vastly different *UMIs/cell*. This strategy may also be useful in other areas, such as the study of rare cell-types. Additionally, the resulting data-set may serve as a benchmark for other methods.

*GTestimate* is available as an open-source R-package (https://www.github.com/Martin-Fahrenberger/GTestimate) and works with all common scRNA-seq data-formats. While *GTestimate’s* default behavior is designed to seamlessly replace *NormalizeData* it is also compatible with a wide variety of other workflows.

## Materials and Methods

### Implementation of *GTestimate*

The user-facing section of our *GTestimate* package was developed in R and handles input and output in the various supported data-formats. The core implementation of the Simple Good-Turing estimator is written in C++ and is heavily based on Aaron Lun’s implementation for the edgeR R-package [23]. This core implementation includes the linear smoothing, which is necessary due to the sparsity of the frequencies of frequencies vector (i.e the frequency of the count values). It further includes a rescaling step which ensures that the estimated relative expression frequencies of all observed genes, plus the sum of probabilities of all unobserved genes (Suppl. Info. 1.1), add up to exactly one [17].

### cta-seq experiment

In the cta-seq experiment we aimed to sequence a selected set of cells from a *typical* scRNA-seq library again at a *ultra-deep* sequencing depth. However, due to sequencing-saturation this quickly becomes prohibitively expensive. We therefore designed a PCR based cell targeted amplification strategy (cta-seq), to selectively amplify all transcripts from a small set of cells, through the use of primers specific to their cell-barcode. This is similar to the TAP-seq protocol [24], which uses gene-specific primers to amplify all transcripts of certain genes.

### Sequencing cta-seq, *typical*

To ensure high quality input material we used leftover cDNA from a previously sequenced sample [25], which had shown high *UMIs/cell* and *genes/cell*. The sample was taken out of -20°C storage and prepared for Illumina sequencing at the Vienna Biocenter Next Generation Sequencing facility using 10X Dual Index Kit TT. We then split the resulting sequencing library into two aliquots and stored the second halve again at -20°C. The first halve was sequenced on a Illumina NovaSeq S4 in paired-end mode with 2×150bp read length and 400 million reads.

### Sequencing cta-seq, *ultra-deep*

Based on the results from the *typical* sequencing run we selected 18 cells of interest for the cta-seq experiment (see below). For these 18 cells we designed PCR primers specific to their cell-barcodes. We then used the second aliquote of the previously prepared sequencing library to perform three rounds of PCR amplification on it using Amplitaq Gold 360 MM (ThermoFisher, cat.: 4398886) supplemented with EvaGreen dye (Biotium, cat.: 31000). We used the following programs in a total volume of 50*µ*l.. PCR1: 1. 95C, 10min; 2. 62C, 30s; 3. 72C, 2min.; 4. Return to 2. x2; 5. 95C, 25s; 6. 62C, 30s; 7. 72C, 2min, fluorescence measurement; 8. 72C, 15s; 9. return to 5. x16. PCR2: 1. 95C, 10min; 2. 62C, 30s; 3. 72C, 2min.; 4. Return to 2. x2; 195C, 25s; 6. 62C, 30s; 7. 72C, 2min, fluorescence measurement; 8. 72C, 15s; 9. return to 5. x16. PCR3: 1. 95C, 10min; 2. 67C, 30s; 3. 72C, 2min.; 4. Return to 2. x2; 5. 95C, 25s; 6. 67C, 30s; 7. 72C, 2min, fluorescence measurement; 8. 72C, 15s; 9. return to 5. x8. Reactions were stopped in step 8 according to fluorescent measurements in log phase. Reaction input in PCRs 2 and 3 were 0.5 *µ*l of the previous reaction. Resulting reactions were purified, and pooled for Illumina sequencing on a NovaSeq S4 in paired-end mode with 2×150bp read length and 400 million reads. The primer sequences used can be found in Suppl. Table 1, PCR1 primers were designed with varying length to achieve similar melting temperatures.

### Data Analysis

All data analysis was performed in R (v4.3.1) using Seurat (v5.0.0) functions at default settings unless stated otherwise.

### Data analysis, cta-seq *typical* depth

We first processed the *typical* depth sequencing data using CellRanger (v7.1.0), this resulted in 20,214 cells. During cell QC we then removed all cells expressing ≤ 1000 or ≥ 5000 genes as well as cells with ≥ 8% mitochondrial reads, with 17,653 cells remaining. We then normalized with Seurat’s *NormalizeData*, selected the top 2000 most variable genes and performed gene-wise z-score scaling. Next we applied PCA and performed unsupervised clustering of cells using the Louvain algorithm [26](resolution = 0.1), based on the first 50 principal components (PCs). This resulted in four cell-type clusters, the smallest cluster (with only 504 cells) was excluded from the subsequent analysis.

From the remaining 17,149 cells we selected 18 cells for targeted amplification, six cells from each of the three remaining clusters. To select a diverse set of cells from each cluster we used the following:

1. We identified the two nearest neighbors for each cell (in PCA space).

2. We excluded cells for which at least one nearest neighbor belonged to a different cluster.

3. For the remaining 16,295 cells, we computed the #UMI-rank, from the number of observed UMIs per cell (ties were broken randomly).

4. Similarly, we computed the 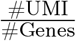 -rank based on the ratio of the number of observed UMIs and the number of observed genes in the cell (ties were broken randomly).

5. Subsequently, we calculated the diversity of each cell and it’s neighbors as the area of the induced triangle of the cell and its neighbors in a #UMI-rank 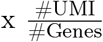-rank plot. The six cells from the two most diverse neighborhoods (i.e. largest triangle area) were selected for amplification.

These steps were designed to cover a diverse set of cells for which the various experimental steps had varying efficiencies. The selection of triplets from the same neighborhoods provided groups of cells with similar gene expression patterns, while the number of UMIs and the number of observed genes were used as proxies for the mRNA capture efficiencies and the health of the isolated cells.

### Data analysis, cta-seq *ultra-deep*

The sequencing data from the *ultra-deep* sequencing run were processed using CellRanger (v7.1.0).

However, due to the high number of PCR cycles during amplification, and the resulting high number of reads for the 18 selected cells, CellRanger’s UMI correction approach was no longer sufficient. Manual inspection of the reads showed that errors in the UMI sequences had inflated the number of unique reads.

This was further exacerbated by a faulty implementation of the UMI-correction approach in the CellRanger software by 10X Genomics. CellRanger erroneously corrects UMIs containing sequencing errors towards other UMIs that also contain sequencing errors. E.g. If we have 3 UMIs: AAAA with 10 reads, AAAT with 2 reads and AATT with 1 read, AATT would be corrected towards AAAT (Hamming Distance 1) and stay as AAAT, eventhough the original 2 AAAT reads would be corrected to AAAA in the same step. We have reported this issue to 10X Genomics on 13th of July 2023, 10X Genomics acknowledge the issue on 14th of July 2023. The issue remains unresolved in CellRanger 7.2.0 (released on the 10th of November, 2023).

To circumvent these issues we extracted the relevant information for each read (count, ensemble gene id, cell-barcode, uncorrected UMI and CellRanger corrected UMI) from the possorted_genome_bam.bam as provided by CellRanger and replicated CellRanger’s read counting workflow in R. As a sanity-check we first used the CellRanger corrected UMIs and achieved the exact same count-matrix as CellRanger. We then used the raw UMIs instead of the CellRanger corrected UMIs, implemented the UMI-tools directional UMI correction approach [27] in R and applied it to correct the UMIs for the 18 selected cells, we then counted again. The resulting count-matrix showed differences for 28% of the non-zero entries when compared to the CellRanger results. We used these improved counts for the *ultra-deep* profiles in all further analysis.

### Comparison of GT and ML using cta-seq

To evaluate the performance of GT and ML based on the cta-seq data-set we estimated the relative gene expression for the 18 selected cells by applying both estimators to the *typical* transcriptomic profiles.

The relative gene expression for the ground-truth *ultra-deep* profiles was estimated with ML. We chose ML to be conservative regarding the performance of GT and since the overestimation due to unobserved genes should be small for the *ultra-deep* profiles Suppl. Figure 9.

### Relative gene expression estimation

We calculated the absolute estimation error for the relative gene expression of the 18 cells by comparing the estimation results of GT and ML based on the *typical* transcriptomic profiles to the ground-truth relative gene expression of the *ultra-deep* profiles. We consider the relative gene expression estimation error of a cell to be the sum of the individual relative gene expression estimation errors in the cell.

### Cell-cell distances

The pairwise Euclidean distances between the 18 cells were calculated in PCA space (as is common for cell-cell distances in scRNA-seq). However, to keep the necessary projections similar to a regular scRNA-seq analysis this space could not simply be constructed based only on the 18 selected cells.

Instead we calculated the projections based on 17,653 cells in the *typical* sequencing run. After normalization there are three pre-processing steps which all depend on the context of a full data-set; Variable gene selection, gene-wise z-score scaling and PCA.

To keep these steps identical for both the GT and ML profiles of the *typical* sequenced cells, as well as the *ultra-deep* profiles we performed them using customized functions. We used the same list of variable genes (calculated based on all 17,653 cells) for the analysis of all profiles. We then scaled the genes in all profiles using the mean and standard deviation of genes calculate based on the full 17,653 cells. Finally we projected all profiles into the same 50 dimensional PCA-space calculated from the full 17,653 cells.

In this PCA-space we calculated the pairwise distances between the ML profiles, between the GT profiles as well as between the ground-truth *ultra-deep* profiles. We then compared the resulting non-zero distances based on GT and ML to the ground-truth *ultra-deep* distances.

### Comparison of GT and ML at different *UMIs/cell*

When analyzing the impact of *UMIs/cell* on the estimation performance we used the cell with the highest number of UMIs after amplification (cell 12, cell-barcode TCTCTGGGTGTGCTTA) and the cell with the second highest number of UMIs after amplification (cell 15, cell-barcode GGCTTTCGTGTGTCGC).

We generated 1000 randomly sampled profiles at each *UMIs/cell* level by drawing genes from the *ultra-deep* count-vector, weighted by count and with replacement. The 20 *UMIs/cell* levels at which we sampled were chosen equidistant in log10-space from 100 to 100,000 (i.e. 100, 143, 206, 297, 428, 615, 885, 1274, 1832, 2636, 3792, 5455, 7847, 11288, 16237, 23357, 33598, 48329, 69519, 100000 *UMIs/cell*). We then applied GT and ML respectively to these sampled profiles to estimate their relative gene expression.

### Relative gene expression estimation

To asses the relative gene expression estimation performance of GT and ML we compared their estimates for each sampled profile from cell 12 to the relative gene expression of the full *ultra-deep* profile of cell 12, and calculated the absolute error.

### Cell-cell distance estimation

To asses cell-cell distance estimation performance we calculated the Euclidean distances between the relative gene expression profiles of pairs of sampled profiles (either from cell 12 twice or from cell 12 and cell 15) based on GT and ML. We calculated the true distance based on the full *ultra-deep* profiles.

### Downstream analysis

#### Data analysis, pbmc3k

The pbmc3k data-set was downloaded from 10X Genomics [19] and processed following Seurat’s “Guided Clustering Tutorial” [28]. In short:

During QC we filtered out genes expressed in less than 3 cells, and cells with less than 200 expressed genes. We then filtered out cells with *>* 5% mitochondrial reads and finally we removed all cells expressing more than 2,500 genes.

During preprocessing cells were normalized using either Seurat’s *NormalizeData* or *GTestimate* at default settings. For both normalization methods individually, we then identified variable genes and z-score scaled the data, followed by calculation of the top 10 PCs. Based on these PCs we then constructed the neighborhood graphs and performed unsupervised Louvain clustering (resolution = 0.5). Finally we calculated the UMAP for both conditions and annotated clusters based on marker gene expression, following the Seurat tutorial.

#### Data analysis, developing pancreas

The pancreas endocrinogenesis day15 dataset was downloaded [29] and imported into R to be processed using Seurat. We only used the spliced counts and normalized them using *GTestimate* and *NormalizeData*; from there on all following steps were performed identically for the two approaches.

First we identified variable genes and performed gene-wise z-score scaling, followed by calculation of the top 50 PCs. Based on the PCs we constructed the neighborhood graph and performed unsupervised Louvain clustering (resolution = 0.4). Finally we calculated the UMAP.

We manually adjusted the cluster numbering (and thereby their color) for Fig. 2c and Fig. 2d. to have consistent cluster-colors from left to right.

#### Data analysis, Spatial Transcriptomics

The stxBrain data-set of sagital mouse brain slices from 10X Genomics was downloaded using the SeuratData R-package. In our analysis we focused on the anterior1 slice of the data-set following Seurat’s “Analysis of spatial datasets (Sequencing-based)” vignette [30].

Our analysis differs from the vignette only in the normalization methods used. While the vignette uses *sctransform*[31] for spot-wise normalization we instead used *NormalizeData* and *GTestimate*. Direct comparison of GT and ML to *sctransform* on the basis of relative gene expression is not possible, since *sctransform* does not calculate relative gene expression levels. Normalization was followed by variable gene selection and gene-wise scaling. We then calculated the first 30 PCs and used them to construct the neighborhood graph, perform unsupervised Louvain clustering and calculate the UMAP.

#### Availability of supporting source code and requirements

1. Project name: GTestimate
2. Project home page: https://github.com/Martin-Fahrenberger/GTestimate
3. Operating system(s): Platform independent
4. Programming language: R, C++
5. Other Requirements: devtools, sparseMatrixStats
6. License: GPL3

*GTestimate* is available as an open-source R-package on github (https://www.github.com/Martin-Fahrenberger/GTestimate). All code for the analysis in this paper, from raw-data to figures, is available on github (https://www.github.com/Martin-Fahrenberger/GTestimate-Paper).

## Supporting information

Supplemental Table 1 - Primer Design

## Data Availability

Processed cta-seq data, such as count-matrices, are available via GEO (https://www.ncbi.nlm.nih.gov/geo/), accession number GSE268930. Due to patient privacy concerns raw sequencing data will be made available through controlled access at the European Genome-Phenome Archive (EGA) upon publication.

### List of Abbreviations

cta-seq: cell targeted PCR-amplification followed by sequencing
GT: Good-Turing estimator
ML: Maximum Likelihood estimator
PC: principal component
scRNA-seq: single-cell RNA-sequencing;

## Authors’ contributions

MF and AvH conceived this project, CE and MF developed cta-seq, CE performed the cta-seq wet-lab experiments, MF implemented GTestimate and analyzed the data. MF wrote the manuscript with input from CE and AvH. All authors read and approved the final version of the manuscript.

## Acknowledgments

We thank Oliver L. Eichmüller for contributing the cDNA-library used in the cta-seq experiment and for his feedback during discussions. We thank all members of CIBIV for their valuable feedback throughout this project. We also thank Thomas Grentzinger from the Vienna BioCenter Core Facilities GmbH (VBCF) Next Generation Sequencing Unit for consultation and sequencing.

## 1 Supplementary Materials

### 1.1 The Missing Mass

Besides improving the relative expression estimates of observed genes, GT can also estimate the sum of the relative frequencies of all unobserved genes. This can be viewed as the probability *p*_0_ that a next hypothetical UMI would be of a currently unobserved gene. We have therefore termed *p*_0_ the missing-mass of the relative gene expression distribution.

The missing-mass for each cell is estimated from the number of genes with a UMI count of one (*N*_1_) and the sum of all counts (Σ _*g*_ *c*_*g*_) as has previously been discussed [16, 17].

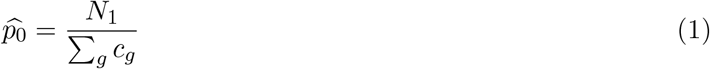

When applied to a Seurat or SingleCellExperiment object in R *GTestimate* saves the estimated 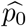 for each cell into a meta-data vector called “missing_mass”.

The Simple Good-Turing estimator scales the relative frequencies (including *p*_0_) to ensure

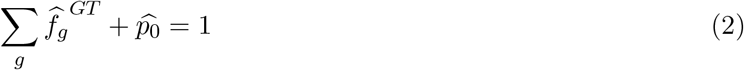

for each cell.

Suppl. Equation 1 provides insight into the amount of information present for each cell, which may warrant further study. E.g. the missing-mass in the cta-seq experiment is substantially reduced after cell targeted amplification of reads (Suppl. Fig. 9).

Due to the typically low *UMIs/cell*, this missing mass of a cell in scRNA-seq can be quite substantial (Suppl. Fig. 10).

### 1.2 Supplementary Tables

**Table 1:** Characteristics of the regression line of the estimated vs. ground-truth distances for the cta-seq data (Fig. 1d).

| Method | Slope | Sum of absolute Residuals | Intercept | Sum of absolute Errors |
| --- | --- | --- | --- | --- |
| ML | 1.529 | 1511.317 | 0.955 | 3258.049 |
| GT | 1.302 | 1263.276 | -0.408 | 2093.645 |

### 1.3 Supplementary Figures

**Figure 1.**
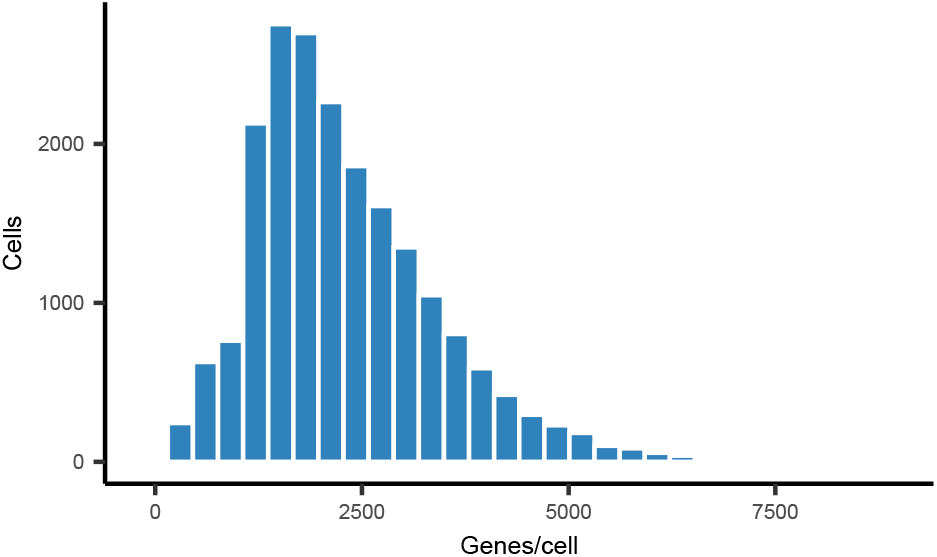
: Histogram showing the number of observed genes per cell for the 17,653 cells in the cta-seq sample before amplification (*typical*).

**Figure 2:**
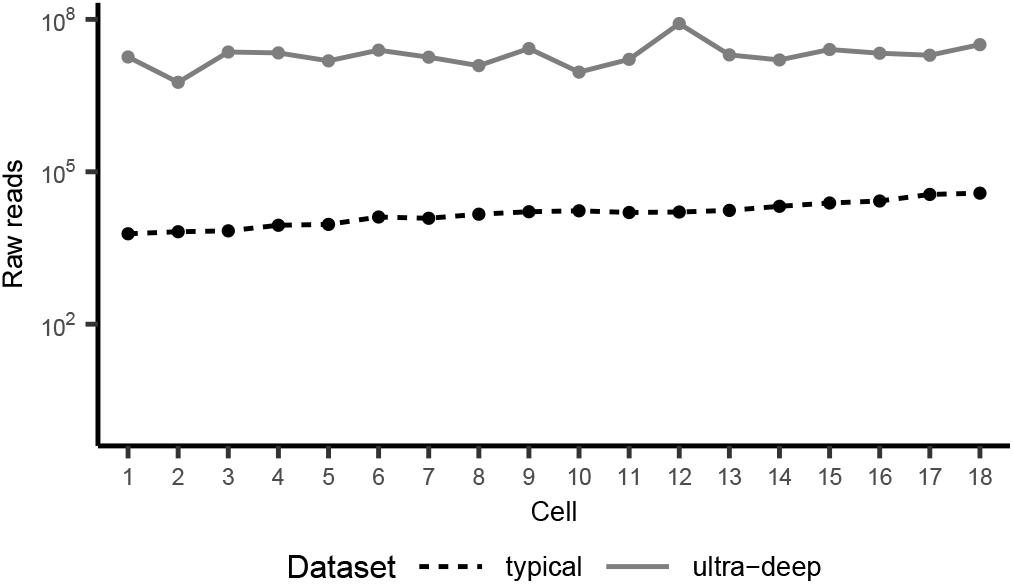
Raw read counts per cell before (*typical*) and after (*ultra-deep*) amplification for the 18 selected cells in the cta-seq experiment.

**Figure 3:**
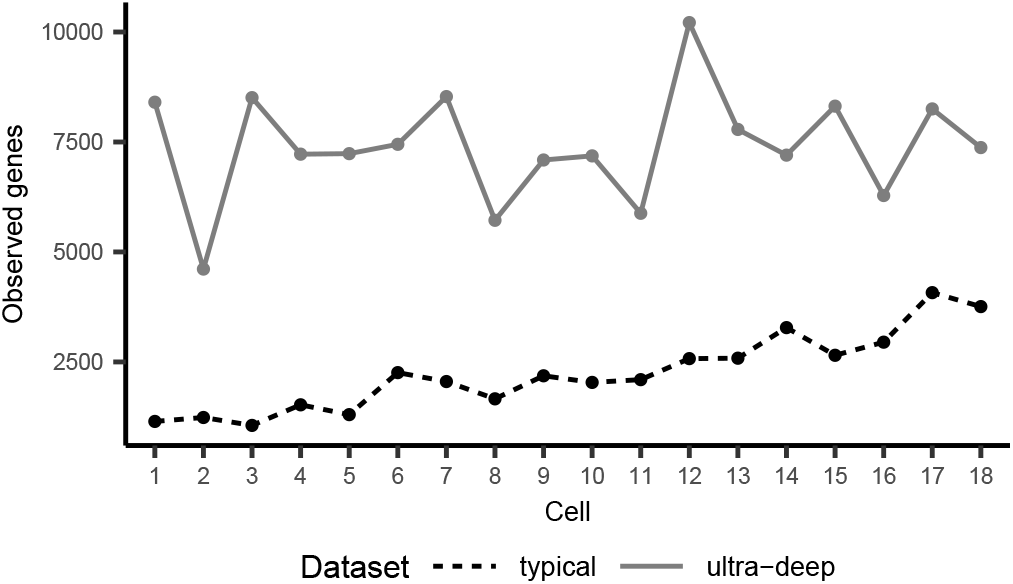
Number of observed genes before (*typical*) and after (*ultra-deep*) amplification for the 18 selected cells in the cta-seq experiment.

**Figure 4:**
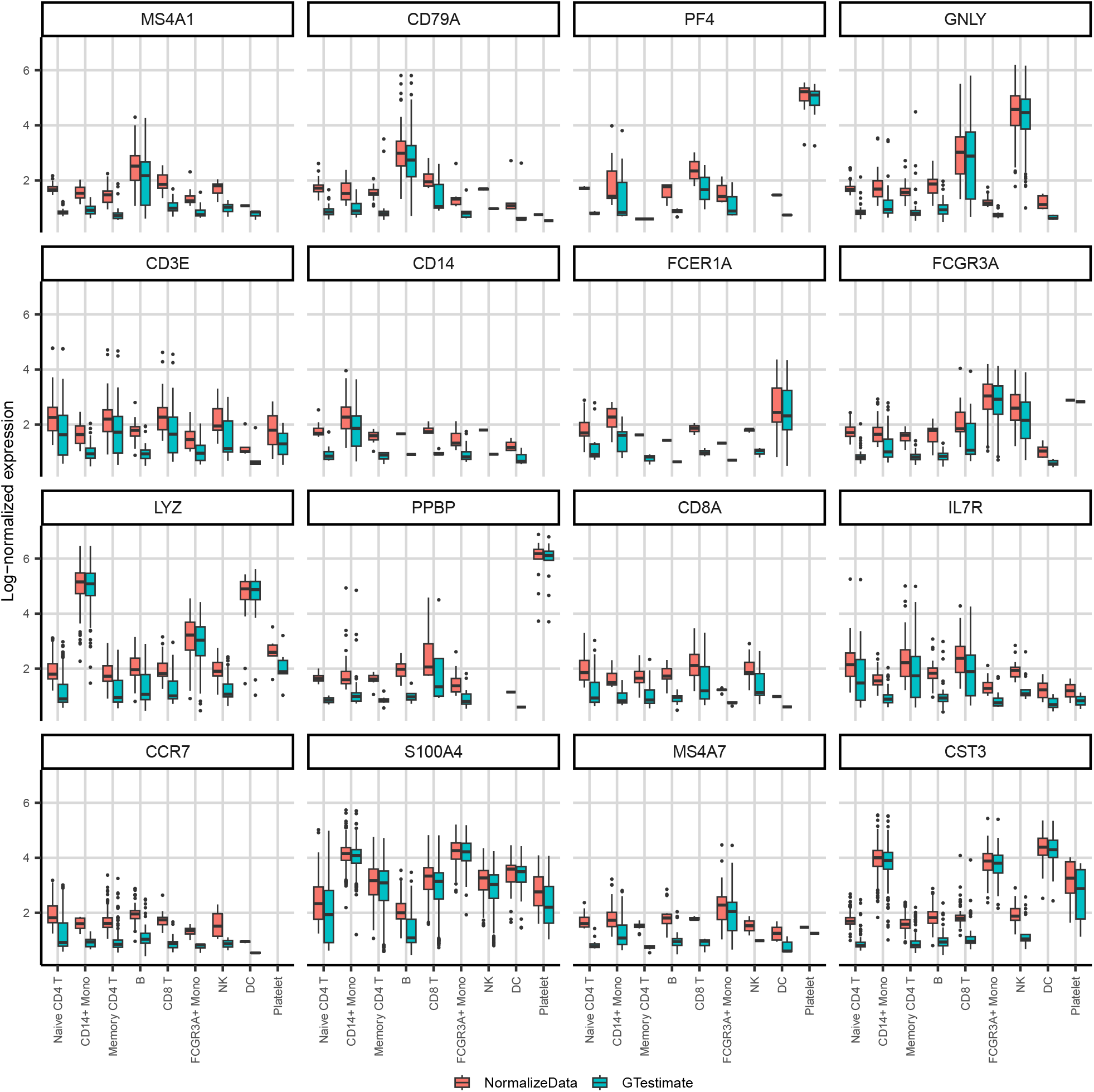
Log-normalized expression of all cell-type markers described in Seurat’s pbmc3k tutorial (zeroes not shown).

**Figure 5:**
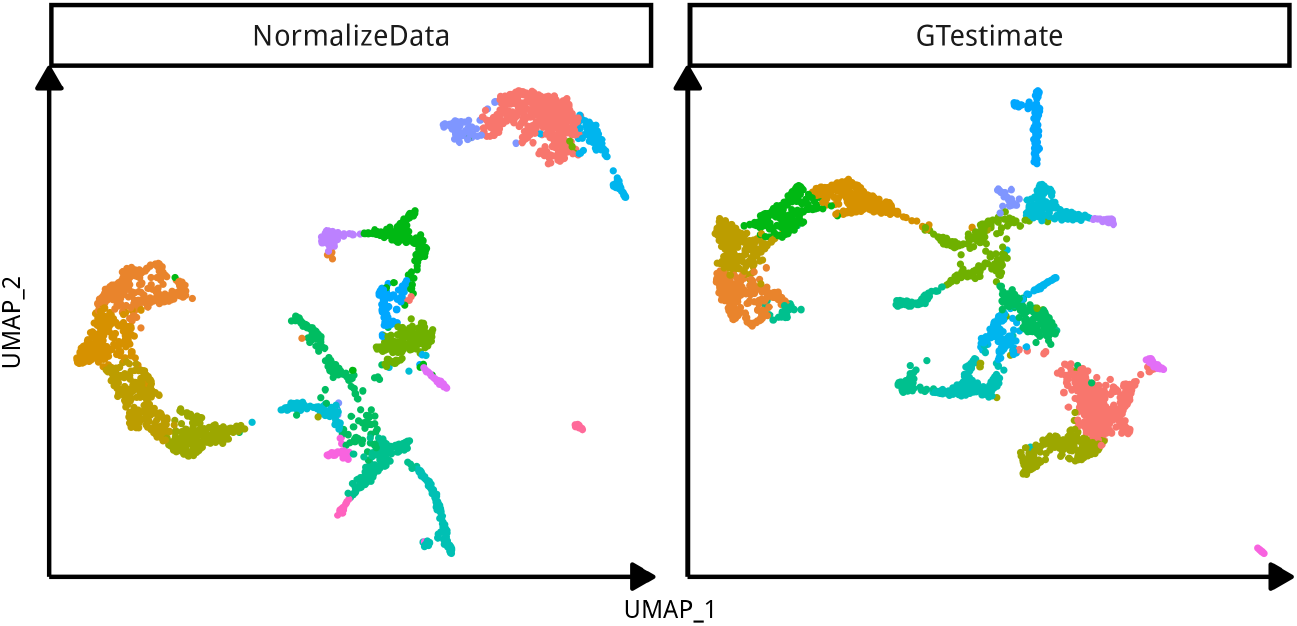
UMAPs visualizing the clustering of Spatial Transcriptomics spots, based on *NormalizeData* **(left)** and *GTestimate* **(right)** for the mouse brain Spatial Transcriptomics data-set.

**Figure 6:**
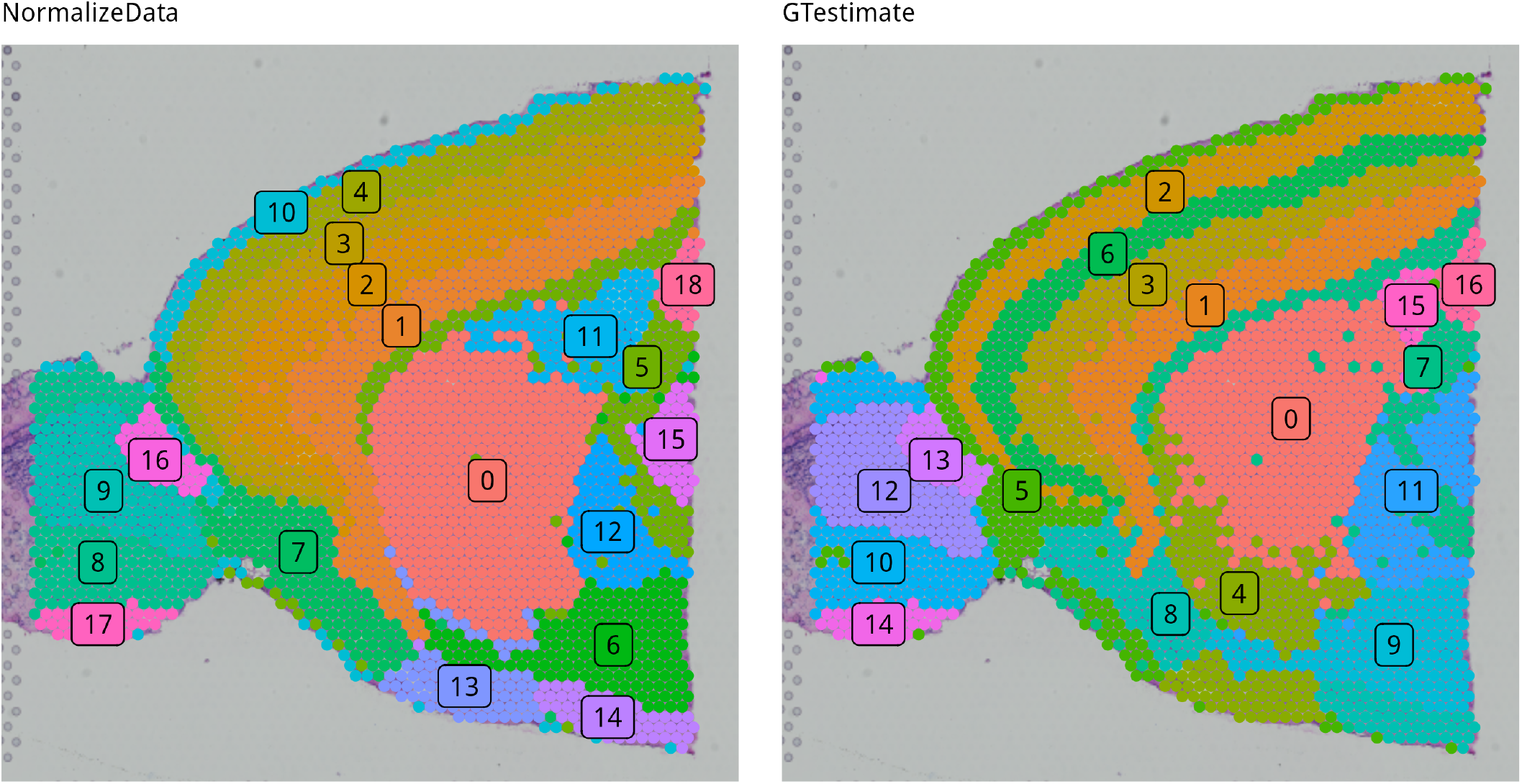
Visualization of the different clusters based on *NormalizeData* **(left)** and *GTestimate* **(right)** for the mouse brain Spatial Transcriptomics data-set.

**Figure 7:**
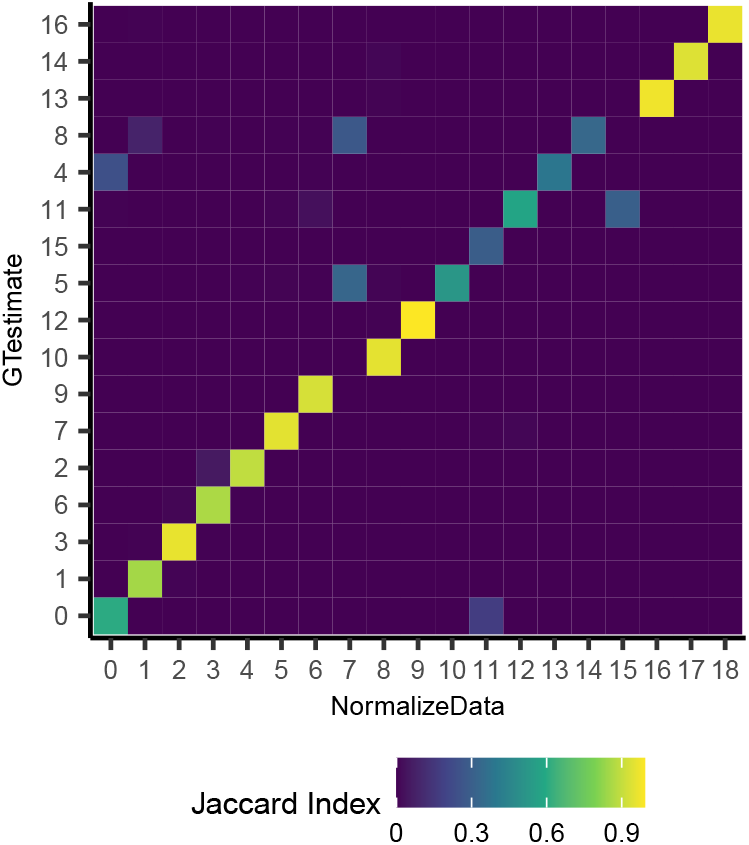
Similarity of the clusters based on *NormalizeData* and *GTestimate* as represented by the Jaccard Index. Clusters on the y-axis have been rearrange to maximize diagonal entries using the Hungarian Algorithm.

**Figure 8:**
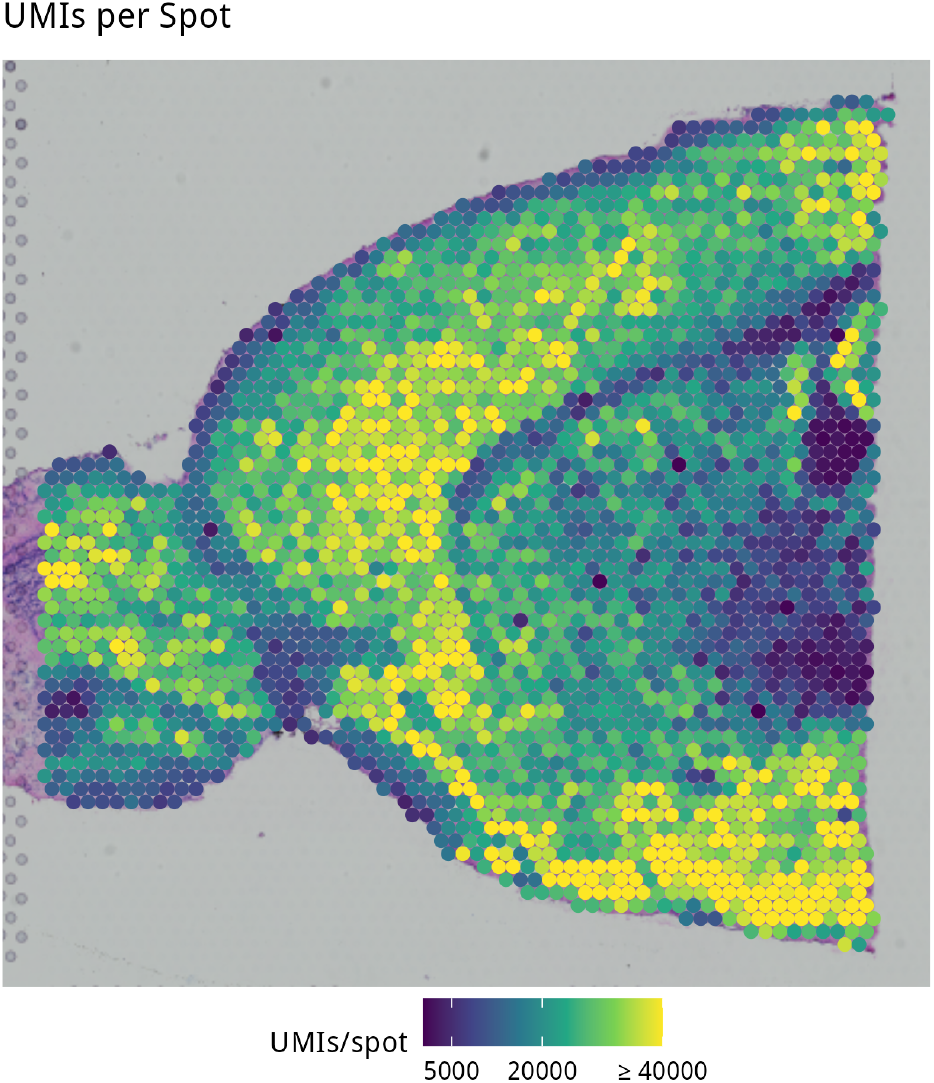
*UMIs/spot* in the Spatial Transcriptomics mouse brain data-set.

**Figure 9:**
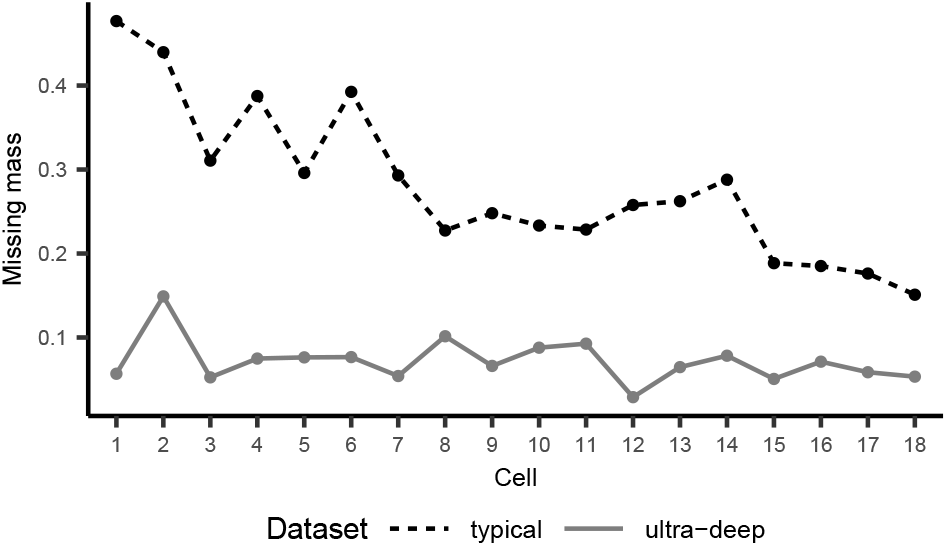
Missing mass before (*typical*) and after (*ultra-deep*) amplification for the 18 selected cells in the cta-seq experiment (see Suppl. Section 1.1).

**Figure 10:**
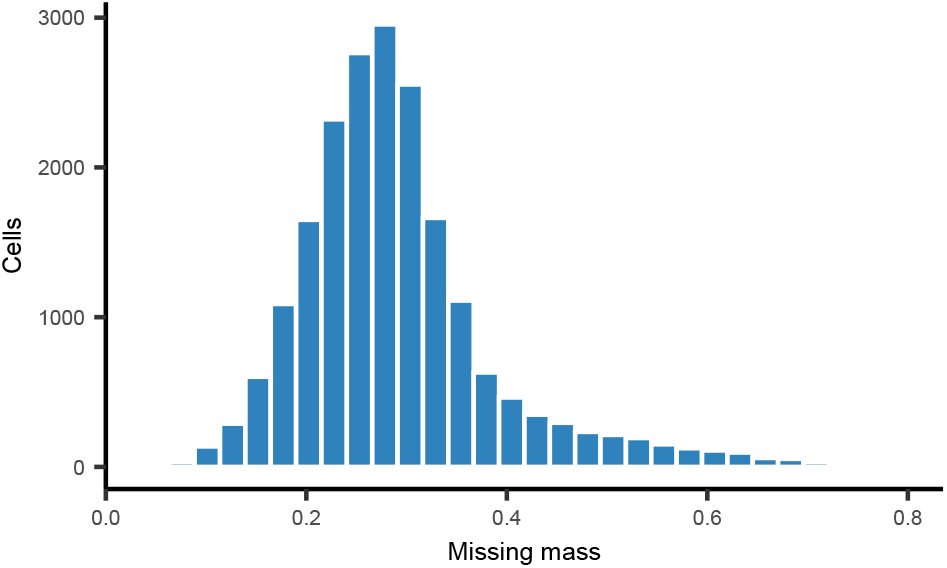
Histogram showing *GTestimate*’s missing mass estimates per cell for the 17,653 cells in the cta-seq sample before amplification (*typical*).

